# Measuring nucleus mechanics within a living multicellular organism: Physical decoupling and attenuated recovery rate are physiological protective mechanisms of the cell nucleus under high mechanical load

**DOI:** 10.1101/2020.02.05.935395

**Authors:** Noam Zuela-Sopilniak, Daniel Bar-Sela, Chayki Charar, Oren Wintner, Yosef Gruenbaum, Amnon Buxboim

## Abstract

Nuclei within cells are constantly subjected to compressive, tensile and shear forces, which regulate nucleoskeletal and cytoskeletal remodeling, activate signaling pathways and direct cell-fate decisions. Multiple rheological methods have been adapted for characterizing the response to applied forces of isolated nuclei and nuclei within intact cells. However, *in vitro* measurements fail to capture the viscoelastic modulation of nuclear stress-strain relationships by the physiological tethering to the surrounding cytoskeleton, extracellular matrix and cells, and tissue-level architectures. Using an equiaxial stretching apparatus, we applied a step stress and measured nucleus deformation dynamics within living *C. elegans* nematodes. Nuclei deformed non-monotonically under constant load. Non-monotonic deformation was conserved across tissues and robust to nucleoskeletal and cytoskeletal perturbations, but it required intact Linker of Nucleoskeleton and Cytoskeleton (LINC) complex attachments. The transition from creep to strain recovery fits a tensile-compressive linear viscoelastic model that is indicative of nucleoskeletal-cytoskeletal decoupling under high load. Ce-lamin (*lmn-1*) knockdown softened the nucleus whereas nematode ageing stiffened equilibrium elasticity and decreased deformation recovery rate. Recovery lasted minutes due to physiological damping of the released mechanical energy thus protecting nuclear integrity and preventing chromatin damage.

## Introduction

The viscoelastic properties of the nucleus are required to protect the genetic material from applied forces(Furusawa *et al.*, 2015) while permitting flexibility to allow constricted cell migration(Denais *et al.*, 2016; Raab *et al.*, 2016; Irianto *et al.*, 2017) and mechanosensitivity.(Athirasala *et al.*, 2017) Externally applied and cell-generated forces propagate along cytoskeletal filaments and are transmitted to the nuclear lamina across Linker of Nucleoskeleton and Cytoskeleton (LINC) complexes. In response to tensile forces that are transmitted to the nuclear lamina across LINC attachments, the nucleus becomes stiffer concomitantly with tyrosine phosphorylation of the nuclear envelope protein Emerin.(Guilluy *et al.*, 2014) Forces pulling on the nuclear lamina are transduced into biochemical cues via tension-suppressed phosphorylation-mediated disassembly and turnover of lamin-A filaments, (Buxboim *et al.*, 2014) which scale with tissue stiffness.(Swift *et al.*, 2013b) Applied forces were also shown to upregulate transcription rate by stretching chromatin, thus acting directly independent of molecular relays. (Tajik *et al.*, 2016)

Multiple complementary rheological methods have been employed for measuring the stress-strain relationship of amphibian and mammalian cell nuclei over a range of deformation length scales and dynamic profiles.(Stephens *et al.*, 2018) Indentation experiments of isolated nuclei(Lherbette *et al.*, 2017) and nuclei within adhering cells (Krause *et al.*, 2013) highlighted the stiffness rendered by condensed chromatin, with contributions from linker DNA and inter-nucleosomal interaction.(Shimamoto *et al.*, 2017) Micromanipulation techniques that typically apply intermediate strains separate between small deformation regime dominated by chromatin and large deformations that are resisted by strain-stiffening of the lamina.(Stephens *et al.*, 2017) Compared with indentation and micromanipulation techniques, micropipette aspiration induces large deformations.(Hochmuth, 2000; Gonzalez-Bermudez *et al.*, 2019) Nuclei responded to applied suction as a viscoelastic solid(Guilak *et al.*, 2000; Rowat *et al.*, 2005) spanning over a broad spectrum of timescales(Dahl *et al.*, 2005) showing decreased deformability with cell differentiation(Pajerowski *et al.*, 2007). Viscous and elastic contributions were affiliated with A- and B-type lamins, respectively.(Swift *et al.*, 2013a)

*In vitro* mechanical characterization of isolated nuclei and nuclei within cells has made fundamental contributions. However, these studies cannot account for the modulating effects of the multiscale physiological surroundings that consists of the cytoskeleton, the cell cortex and membrane, the extracellular environment of cells and matrix, and tissue architectures. Already within intact cells, nucleus stiffening was shown to increase cytoskeletal strain by serving as a stress concentrator.(Heo *et al.*, 2016). In turn, the physical tethering with the cytoskeleton effectively stiffens the nucleus and slows down the release rate of the stored mechanical energy.(Wang *et al.*, 2018)

To measure the strain-stress relationship of the nucleus within its physiological multicellular settings, we performed creep test of living *C. elegans* nematodes. Nematodes were placed in between two membranes, stretched equiaxially and nucleus deformation dynamics was recorded. Counter to the whole nematodes and cells within tissue, nuclei exhibited a non-monotonic response under constant load. Once critical strain was reached, the nucleus transitioned from creep to strain recovery. This anomalous response was conserved across tissues, cytoskeletal perturbations and nuclear perturbations, yet it required intact LINC attachments. We propose a linear viscoelastic model that superimposes apical compression and tensile stretching which is relaxed due to nuclear decoupling via LINC complex detachment at high load. *Lmn-1* knockdown increased dissipative recovery time and softened nucleus resistance to continuous stresses but had no effect on instantaneous elasticity. Despite maintaining constant Ce-lamin protein level, nuclei became stiffer and deformation recovery became slower with ageing. Nuclei responded within minutes – much slower than *in vitro* recovery times scales over seconds. Hence, the physiological surroundings of the cytoskeleton, extracellular matrix and cells, and tissue architectures attenuate the release rate of the mechanical energy stored within the nucleus thus protecting the nuclear envelope and the genetic material from mechanical damage.

## Results

### Creep test of live nematodes reveals non-monotonic nuclear response to applied stresses

Measuring nucleus mechanics within live nematodes was performed using a custom made device constructed based on a former cell stretcher design (Lammerding and Lee, 2009) and mounted onto an inverted epifluorescence microscope (Fig. 1a). Live nematodes were placed in between two parallel silicon membranes and immersed in M9 buffer. Pulling the two parallel membranes downwards against a static cylinder generated an in-plane uniform and homogenous equibiaxial tensile strain as calculated by tracking the relative displacements of distinct defects. The deformation dynamics of the nematodes was recorded before and during membrane stretching by time-lapse phase contrast and fluorescence imaging (Fig. 1b). Similarly, the deformation dynamics of nuclei within different tissues was evaluated by time-lapse recording of a GFP-fused Ce-lamin fluorescence signal (Fig. 1c).

**Figure-1.**
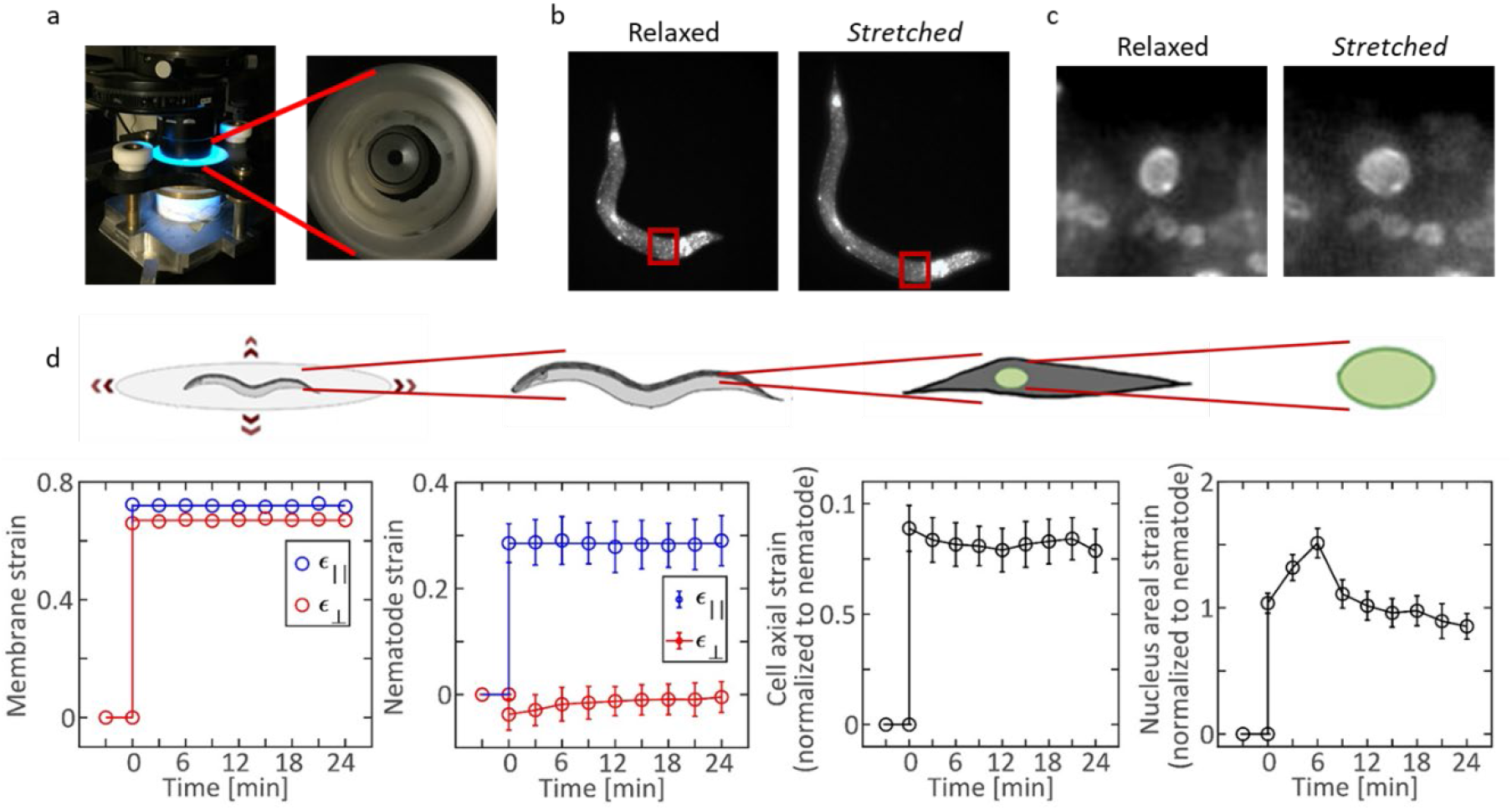
Mechanical creep test of cell nuclei within live nematodes reveals anomalous response to applied stresses. **a**, Live *C. elegans* nematodes are placed between two elastic silicon membranes on a stretching apparatus mounted onto an inverted microscope. **b**, Fluorescent time-lapse images of the nematodes are recorded as the membranes are being radially stretched. **c**, Zoom-in images (red frames in b) of GFP-conjugated Ce-lamin nuclei show induced deformation along the primary axis of the nematode. **d**, Creep test is performed by maintaining a 70% radial strain of the membranes (left). Average dynamic strain profiles of the nematodes (n=3), cells (estimated as the distance between proximal nuclei, n=7) and nuclei (n=12) are evaluated based on time lapse imaging. Unlike nematodes and cells that exhibit a constant elastic axial deformation, nuclei respond non-monotonically to applied stress (right).

We characterized the mechanical response of worms, cells and nuclei to applied stresses using creep test rheology. Specifically, we pulled the membranes at ~70% equibiaxial strain and maintained stretching for 24 min (Fig. 1d-i). As a result, the nematodes were stretched longitudinally but not perpendicular to the main axis of the worm due to their thin cylindrical geometry (Fig. 1d-ii). The nematodes responded elastically to applied stress: instantaneous deformation at t=0, which was maintained as long as the membranes were held stretched. To evaluate the axial deformation of cells, we estimated the distances between the centers of consecutive pairs of segmented nuclei within the obliquely striated myofibrillar lattice of the body wall muscle tissue (Gettner *et al.*, 1995). Nuclear areal strain dynamics was also evaluated using the segmented GFP-signal regions of interest (ROI’s). Both cell and nucleus strains were normalized by worm strains to compensate for differences in the forces that were applied within different nematodes. We found that cells responded elastically to applied stresses just like the nematodes (Fig. 1d-iii). However, nuclei exhibited a complex dynamic response that was marked by a non-monotonic deformation under constant load (Fig. 1d-iv).

### Nucleus mechanics fit a tensile-compressive linear viscoelastic model

The irregular deformation dynamics of nuclei was observed both in muscle and in hypodermis, indicating that it is an invariant property across tissues (Figs. 3a-i,ii), which is a property of individual nuclei in both tissues (Fig. S1). It consists of a creep phase between t=0 and a critical time *t*_*cr*_, where strain increases with time, and a recovery phase at *t* > *t*_*cr*_, where strain decreases (Fig. 2a). To obtain insight into the mechanisms that give rise to the creep-recovery response to load, we tested the effects of the LINC complex on this response. LINC complexes are required for mediating the transmission of forces that are propagated along cytoskeletal filaments to the nuclear matrix of lamins and chromatin and back.(Buxboim *et al.*, 2010) To this end, we performed creep test as described above on Unc-84-null nematodes that lack integral LINC attachments.(Malone *et al.*, 1999; Bone *et al.*, 2014) Remarkably, the irregular deformation dynamics that was conserved across tissues vanished and instead Unc-84-null nuclei responded elastically to applied forces (Fig. 3b).

**Figure-2.**
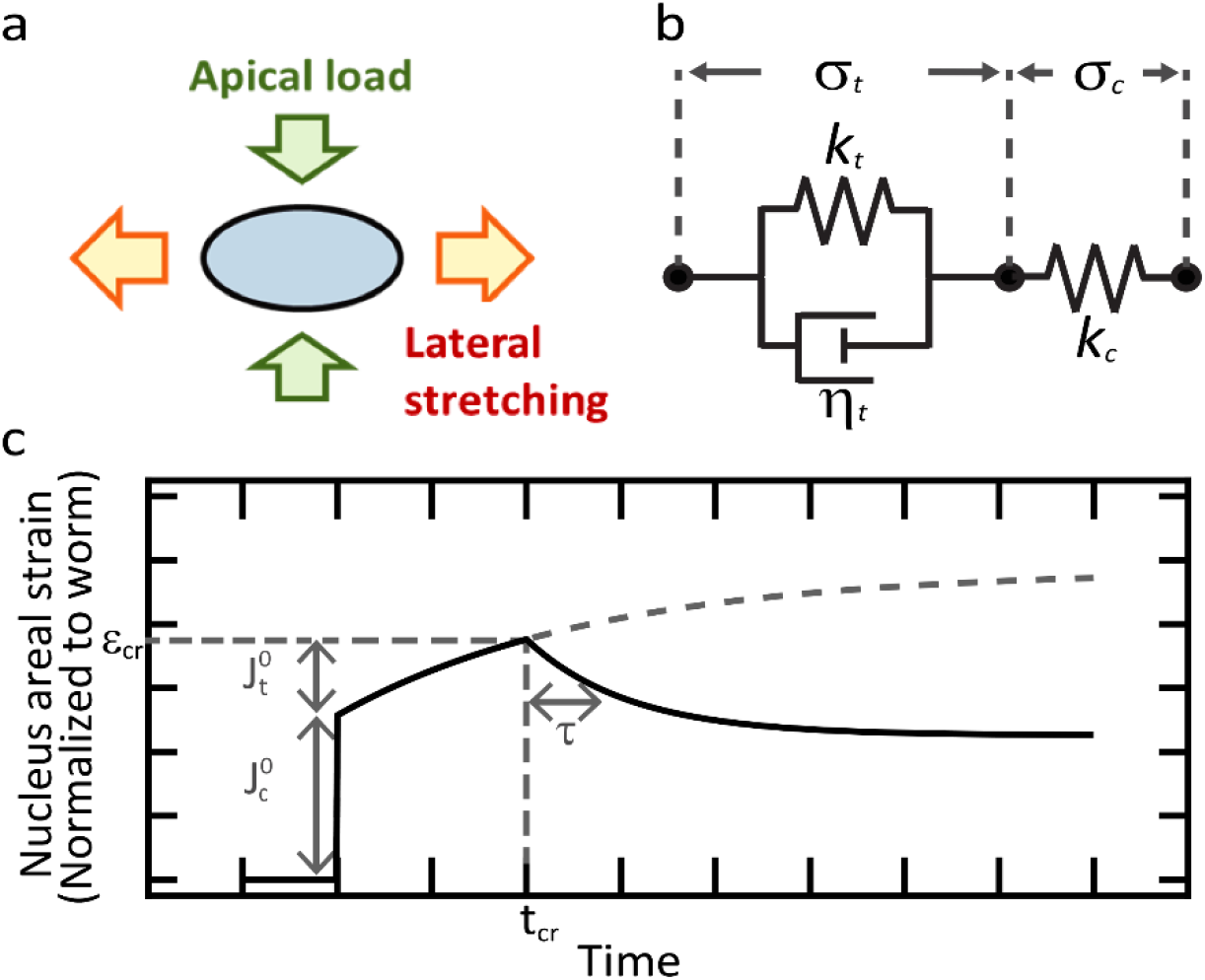
Tensile-compressive viscoelastic model of the cell nucleus in live nematodes. **a**, Nucleus response to constant load consists of an instantaneous elastic response to compressive stresses (*ε*_*c*_), followed by creep response to tensile stresses and partial deformation recovery (*ε*_*c*_). The non-monotonic transition from creep to recovery occurs at critical time *t*_*cr*_ once critical strain *ε*_*cr*_ is reached. **b**, Equiaxial stretching of the two membranes generates tensile stresses on the nuclei that are mediated via LINC attachments as well as compressive stresses that are independent of intact LINC as the membranes are pulled against each other due to in-plane tension. **c**, A minimal viscoelastic model of the nucleus consists of a viscoelastic solid-like Kelvin-Voigt module and an elastic spring connected in series. The Kelvin-Voigt module resists tensile stresses whereas the spring models the response to compressive stresses.

**Figure-3.**
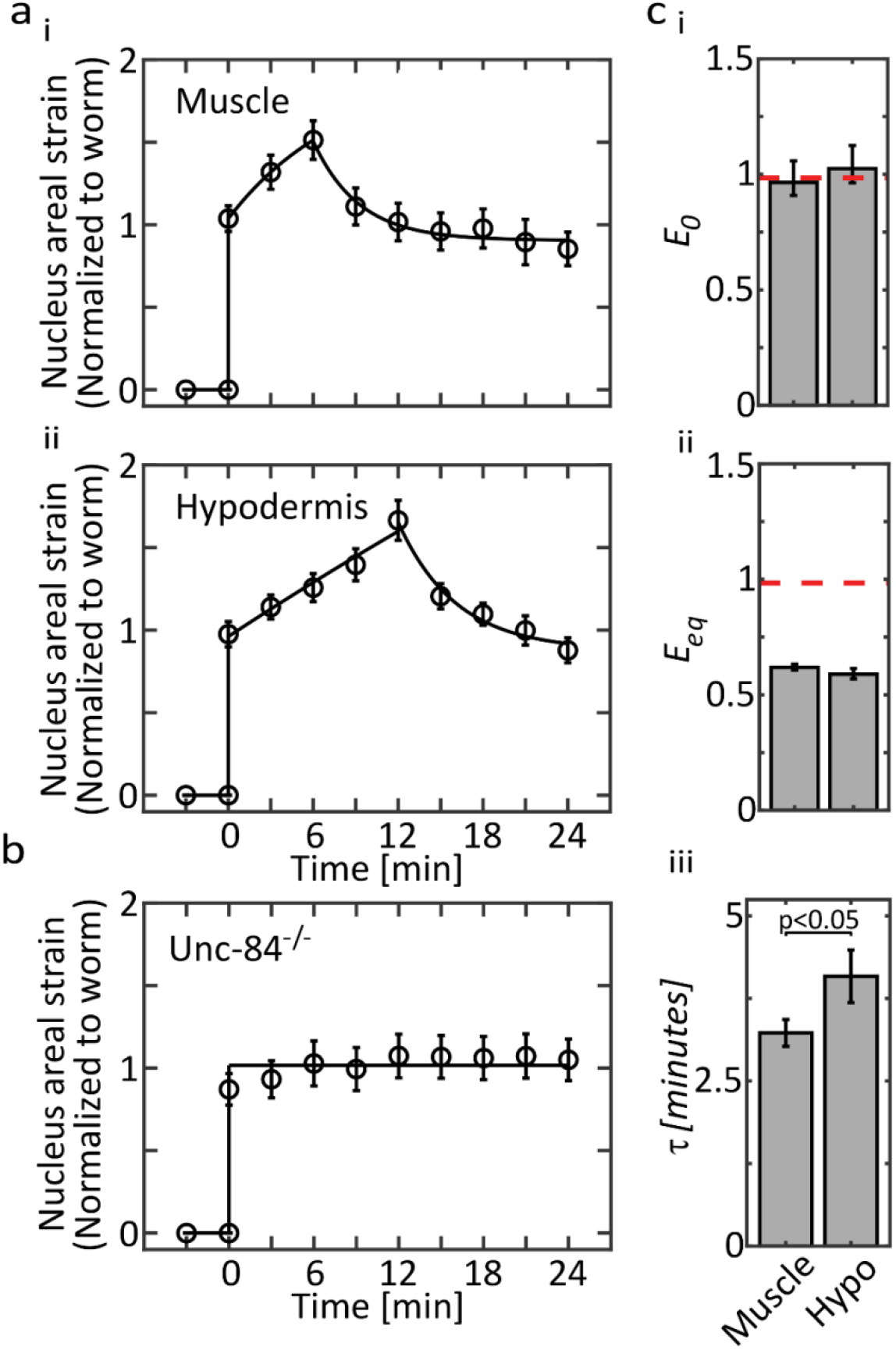
Non-monotonic nucleus deformation is conserved across tissues and depends on LINC attachments. **a**, Creep tests of nuclei in (i) muscle tissue (12 nuclei in 2 nematodes) and in (ii) hypodermis tissue (11 nuclei in 2 nematodes) share a non-monotonic response to applied load. **b**, Nuclei in Unc-84-null nematodes (18 nuclei in 4 nematodes) maintain a constant deformation under a constant load. **c**, Evaluation of the (i) instantaneous and (ii) equilibrium elastic moduli and the (iii) viscoelastic response time. Muscle and hypodermis parameters were evaluated by curve fitting using the tensile-compressive model. Red lines represents *unc-84 null* stiffness evaluated for a purely elastic response.

Based on our nematode stretching experiments, we hypothesize that the LINC complexes undergo physical detachment once a critical strain is reached, leading to nucleoskeletal-cytoskeletal decoupling (NCD). Importantly, even if only a few LINC complexes break, the average loading per attachment will increase on the remaining LINC complexes, thus leading to rapid avalanching detachment of all LINC complexes as reflected here by the sudden decrease in nuclear strain. Despite the fact that NCD occurred at different times in muscle and hypodermis nuclei, it was observed at similar levels of critical strain (Fig. 3a-i,ii). NCD is a protective mechanism of the chromatin from physical tears and breaks due to high tensile strain as was recently reported for a culture model of mammalian cells (Gilbert *et al.*, 2019). In addition to tensile stresses, our membrane stretching apparatus also applies compressive forces on the nuclei as the membranes are pulled downwards and against each other (Fig. 2b). Unlike tensile stresses, compressive load is applied to the nucleus independent of the integrity of the LINC complexes and is maintained constant independent of NCD.

We employ the Kelvin representation standard linear solid (SLS) to model nucleus response to tensile stress *σ*_*t*_ and compressive stress *σ*_*c*_ in living nematodes (Fig. 2c). Nucleus resistance to tensile stresses is modeled by a solid-like Kelvin-Voigt viscoelastic module, which is composed of a spring and dashpot connected in parallel (*k*_*t*_ and *μ*_*t*_). Nucleus resistance to compressive stresses is modeled using a spring (*k*_*c*_). During creep phase, the LINC complexes remain intact and tensile and compressive deformations are superimposed. The strain-stress relationship is:

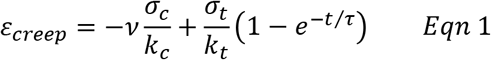

where *τ* = *μ*_*lat*_/*k*_*lat*_ is the Kelvin-Voigt response time and *v* is the Poisson’s ratio of the nucleus responsible for converting compressive loading to lateral deformation. Upon NCD, only compressive stress remains leading to a gradual recovery of the Kelvin-Voigt module with time. In this case nucleus strain asymptotically approaches the instantaneous deformation as imposed by compressive stress:

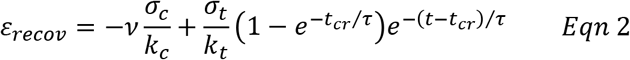

The instantaneous elastic modulus *E*_0_ is evaluated based on creep phase strain at t=0:

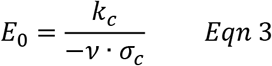

Nuclear response time *τ* and equilibrium elastic modulus *E*_*eq*_ are evaluated by fitting the recovery phase strain dynamics:

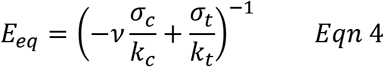

*E*_*eq*_ accounts for steady state elastic resistance to applied stresses of the two springs *E*_*t*_ and *E*_*c*_ in series.

### Muscle and hypodermis nuclei have similar stiffness but different viscosities

Nuclear viscoelastic properties in muscle and in hypodermis tissues were obtained by fitting the average strain dynamics curves (Fig. 3a-i,ii, solid lines). Fitting quality was scored by R-square 0.97 in muscle and 0.98 in hypodermis. Since the nucleus is physically decupled from the surrounding cytoskeleton in Unc-84^−/−^ nematodes, areal strain was fitted using an elastic spring (R-square 0.93, Fig. 3b, solid line). The instantaneous elastic modulus, equilibrium elastic modulus and recovery response time were evaluated for muscle, hypodermis and Unc-84^−/−^ nuclei based on fitting of the normalized nuclear strain curves. The instantaneous and equilibrium elastic moduli, *E*_0_ and *E*_*eq*_, were both equal in muscle and hypodermis nuclei (Fig. 3c-i,ii). Since instantaneous deformation in our stretching experiments is dominated by compressive stresses, *E*_0_ is equal in Unc-84 null and LINC-intact nuclei (Fig. 3c-i). However, *E*_*eq*_ is higher because tensile stresses are disrupted only in Unc-84 null nuclei whereas *E*_*eq*_ in LINC-intact nuclei is set by *k*_0_ and *k*_*eq*_ springs in series (Fig. 3c-ii). Response time *τ* was relatively short in muscle nuclei and long in hypodermis nuclei (Fig. 3c-iii). Taken together, our stretching measurements of nuclear mechanics in live nematodes support the proposed linear viscoelastic model that accounts for NCD at critical strain and combines viscoelastic tensile response and elastic compressive response (Fig. 2c).

### Chromatin and lamin control the viscoelastic properties of the nucleus

To obtain a mechanistic understanding of nuclear mechanics, we studied how nucleus resistance to applied stresses depends on cytoskeletal and nucleoskeletal organization in live nematodes. We perturbed filamentous actin polymerization by treating live nematodes with Latrunculin-A (Lat-A; Fig. 4a-i) and disrupted the LINC complex attachments by feeding the nematodes with RNAi against *unc-84* (unc84i; Fig. 4a-ii). In addition, we targeted the nuclear lamina and chromatin by feeding live nematodes with RNAi against *lmn-1* (lmn1i, Fig. 4a-iii) and treatment with Trichostatin-A (TSA, 4a-iv), respectively. TSA is a potent Class-I and −II histone deacetylase inhibitor (HDACi) that is broadly used for chromatin de-condensation. This is achieved by rapid induction of a global increase in histone acetylation, thus removing positive charges and relaxing DNA interactions. Acetylation of histone H4K16 that interferes with the structural compaction of the 30 nm chromatin fibers was also reported.(Shogren-Knaak *et al.*, 2006; Robinson *et al.*, 2008; Chen and Li, 2010)

**Figure-4.**
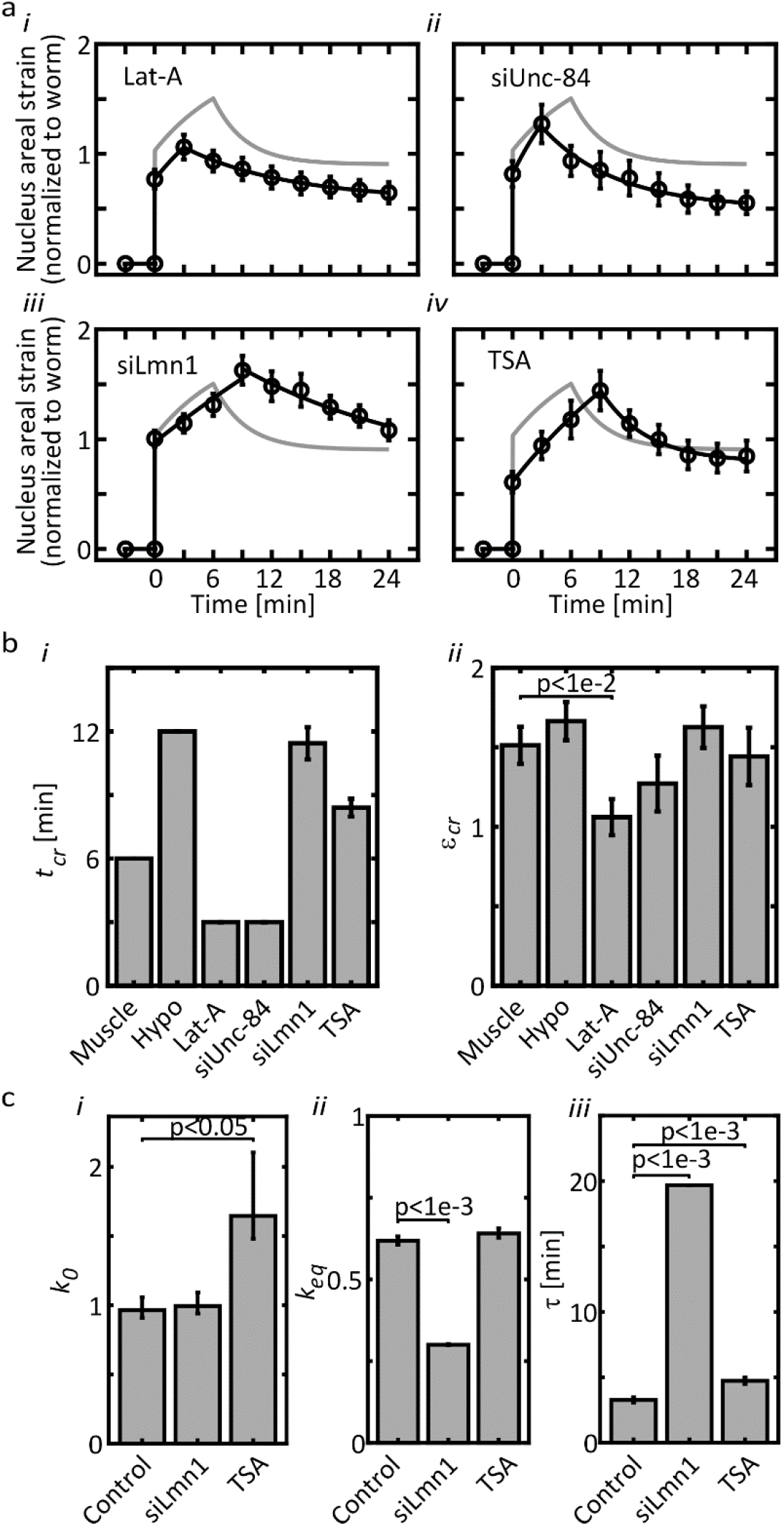
Cytoskeletal and nuclear perturbations modulate nucleus deformation dynamics in accordance with the tensile-compressive viscoelastic model. **a**, Creep tests of muscle tissue nuclei in nematodes treated with (i) Lat-A (3 nematodes, 14 nuclei), (ii) siUnc-84 (3 nematodes, 14 nuclei), (iii) siLmn-1 (4 nematodes, 16 nuclei) and (iv) TSA (3 nematodes, 10 nuclei) are fitted using the tensile-compressive model. Non-treated nucleus deformation curves are plotted in gray as reference. **b**, Evaluation of the (i) instantaneous and (ii) the equilibrium elastic moduli and (iii) viscoelastic response time. Lat-A: Latrunculin-A. RNAi: RNA interference. TSA: Trichostatin-A.

Changes to the viscoelastic properties of the nucleus due to the cytoskeletal and nucleoskeletal perturbations were evaluated via creep test and model fitting as described above. Compared with control nuclei (Fig. 4; light gray), both actin de-polymerization and LINC disruption led to an overall decrease in nucleus compliance (Fig. 4a-i,ii). Critical strain for NCD appears to be a conserved property across tissues and invariant to nuclear perturbations. However, both cytoskeletal perturbations decreased critical time and strain compared with untreated muscle control (Fig. 4b-i,ii). Relative to non-perturbed muscle nuclei, *Lmn-1* knockdown led to a shallower creep slope and a higher critical strain (Fig. 4a-iii). Model fits of nucleus perturbations indicate that knockdown of *Lmn-1*, which in the nematode combines properties of both A- and B-type mammalian lamins,(Liu *et al.*, 2000; Lyakhovetsky and Gruenbaum, 2014) softened the equilibrium elastic modulus twofold and prolonged response time sevenfold (Figs. 4c-ii,iii). Increase in *τ* is consistent with a decrease in the restoring force due to softening of *k*_*t*_ (Fig. 2c), in tune with previously reported lamin-A knockdown experiments in mammalian cells (Lammerding *et al.*, 2006) and our unpublished data. Instantaneous elasticity was dominated by chromatin rather than Ce-lamin (Fig. 4c-i). On the other hand, chromatin de-condensation attenuated instantaneous deformation in response to impact (Fig. 4a-iv) in a manner that effectively corresponds to stiffening of *E*_0_ by > 60% (Fig. 4c-i). However, cytoskeletal effects of TSA should be considered (see discussion).

### Ageing alters long term stiffness without significantly decreasing normalized lamin levels

*C. elegans* is an established model organism in the research of ageing and associated laminopathic genetic disorders.(Haithcock *et al.*, 2005) To study how nuclear mechanics changes with ageing, we performed creep test of L4 larvae (D0), two-day old (D2) and four-day old (D4) adult nematodes (Fig. 5a). We first compared Ce-lamin protein levels between D0, D2 and D4 nematodes via western blotting of whole nematodes lysates (Fig 5b-i). Since D0 nematodes are significantly smaller despite having an equal number of cells and nuclei as D2 and D4 nematodes (Fig. 5a), we normalized Ce-lamin intensities by (total protein)/(nematode volume). Based on three biological replicas, we excluded significant differences in Ce-lamin protein levels with ageing (Fig. 5b-ii).

**Figure-5.**
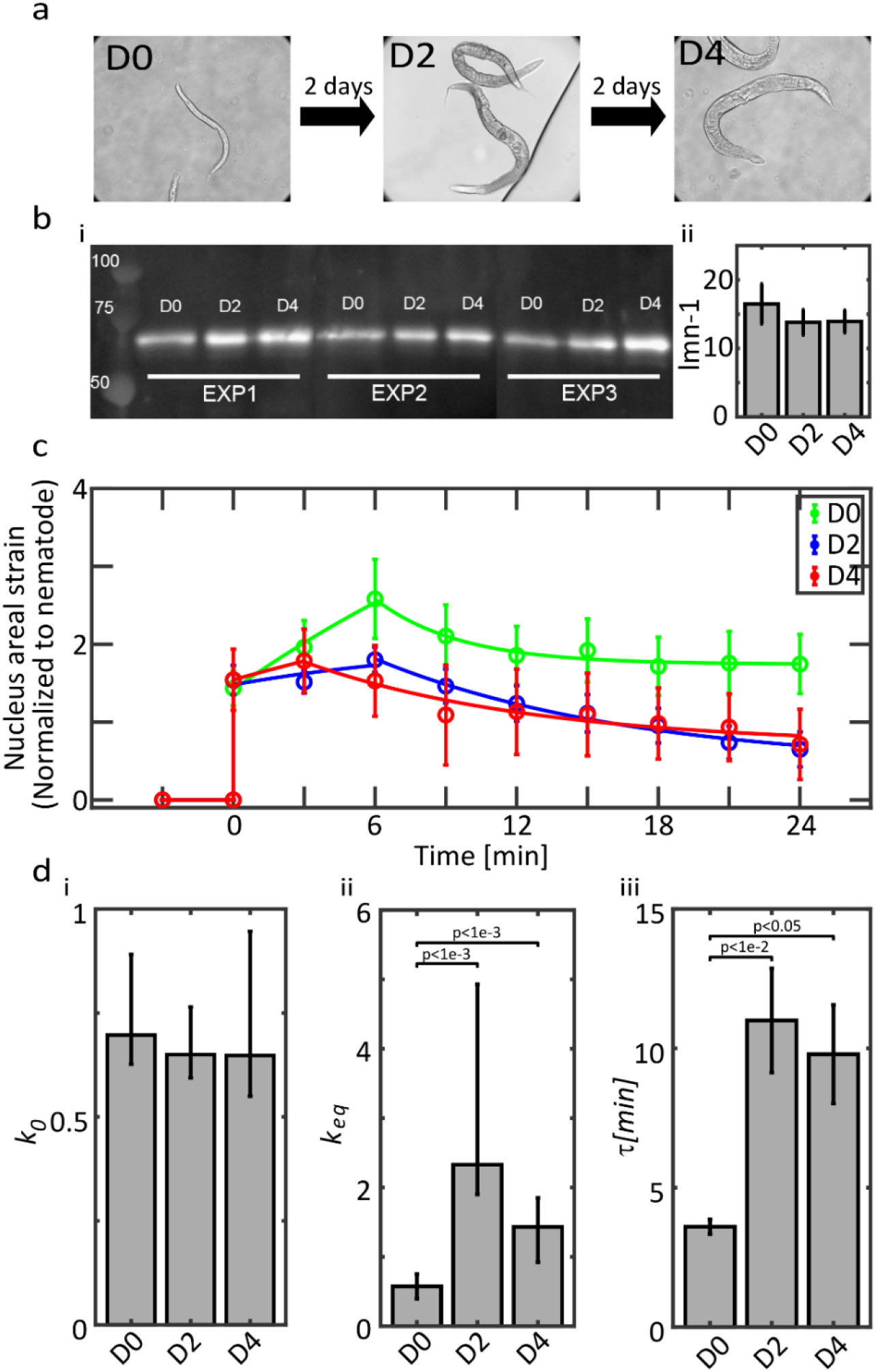
Ageing alters nucleus elasticity and viscosity. **a**, *C. elegans* nematodes grow in size as they age. **b**, (i) Western blot of *C. elegans* samples, stained for Ce-lamin (ii) Ce-lamin levels normalized according to total protein count and nematode size. **c**, Nucleus strain dynamics of Day-0 (3 nematodes, 10 nuclei), Day-2 (3 nematodes, 9 nuclei), and Day-4 (2 nematodes, 5 nuclei) nematodes. **d**, Evaluation of the (i) instantaneous and (ii) the equilibrium elastic moduli and (iii) viscoelastic response time.

Instantaneous strain was comparable between D0, D2 and D4 creep test measurements (Fig. 5c). However, strain dynamics slope during creep phase was high in D0 nematodes and low in aged D2 and D4 nematodes. With ageing, critical strain decreased between D0 and D2 and critical time was shorter in D4 nematodes, thus recapitulating the cytoskeletal effects that we observed following actin de-polymerization and LINC disruption (Fig. 4b-i,ii). Instantaneous stiffness was invariant to nematode age. However equilibrium stiffness was low in D0 nuclei and high in D2 and D4 nuclei (Fig. 5c-i,ii) and response time was threefold longer in aged nuclei (Fig. 5c-iii). Collectively, we find that ageing increases the elastic resistance of nuclei to applied stresses and the dissipative response time required for reaching mechanical equilibrium.

## Discussion

Here we present, to the best of our knowledge, the first nucleus rheology study in a live multicellular organism. Using a custom designed worm stretching device, we performed a creep test and identified anomalous deformation dynamics of individual nuclei. The non-monotonic transition from creep to recovery is indicative of NCD once critical strain is reached, as experimentally supported by the elastic deformation of SUN-domain Unc-84 knockout nuclei that lack intact LINC complex attachments. Nucleus mechanics is well modeled by a three-elements linear viscoelastic material (SLS model) (Guilak *et al.*, 2000) that superimposes LINC-dependent tensile stresses and LINC-independent compressive stresses. Disruption of filamentous actin polymerization and knockdown of *unc-84* do not ablate all LINC complexes but rather decrease their number. Hence, earlier NCD at lower critical strain is likely attributed to the increase in the average load per LINC attachment (Fig. 4b-i,ii). The anomalous response of the nucleus to applied stresses is conserved across tissues and across conditions. LINC complex repair appears to be hindered for approximately 20 minutes following NCD, optionally due to retraction of the cytoskeletal filaments away from the nucleus. Tracking LINC repair during longer time periods following NCD was impeded due to loss of nematode viability during creep test.

Chromatin de-condensation by HDACi was reported to either soften nucleus resistance to small deformations (Chalut *et al.*, 2012; Krause *et al.*, 2013; Haase *et al.*, 2016; Shimamoto *et al.*, 2017) or not to stiffen nucleus resistance to large deformations.(Stephens *et al.*, 2017) Counter to these *in vitro* findings that apply forces directly on the nucleus, here we probe nucleus deformation dynamics while indirectly stretching the nuclei by pulling on the entire nematodes (Fig. 1). Hence, we cannot exclude cytoskeletal effects of HDACi. TSA was shown to inhibit HDAC6, which acts on α-tubulin both *in vivo* and *in vitro*.(Matsuyama *et al.*, 2002) In this manner, TSA can promote acetylation of α-tubulin, likely at the *ε*-amino group of the N-terminus conserved lysine residue (Lys40) (MacRae, 1997; Rosenbaum, 2000), thus contributing to the stabilization of the cytoplasmic microtubule filaments.(Matsuyama *et al.*, 2002) In particular, TSA is expected to stiffen skeletal muscle cells that are rich in microtubule networks.(Robison and Prosser, 2017). We therefore conclude that the decrease in instantaneous nuclear strain due to TSA treatment cannot be definitively attributed to chromatin de-condensation. In fact, it may reflect altered muscle cell and tissue level mechanics rather than nuclear stiffening due to chromatin de-condensation (Fig. 4b-i).

*C. elegans* nuclei exhibit ageing-related modifications that include changes to nuclear morphology and loss of peripheral heterochromatin, which are also observed in accelerated ageing disorders.(Haithcock *et al.*, 2005) Here we performed creep test of L4, and D2 and D4 adult nematodes (Fig. 5a) to characterize the effects of nematode age on nucleus mechanics (Fig. 5c). We found that in aged nematodes NCD was observed earlier and at lower critical strain compared with young nematodes (Fig. 5c). A decrease in critical strain is consistent with equilibrium nucleus stiffening and prolonged response time (Fig. 5d-ii,iii). However, nuclei maintain constant instantaneous elasticity with age, which renders mechanical strength to the nucleus against impact (Fig. 5d-i). Our *in vivo* measurements reproduce *in vitro* studies that report stiffening of equilibrium elasticity, which is a hallmark of HGPS as well as normal ageing in amphibian and mammalian cells.(Dahl *et al.*, 2006; Verstraeten *et al.*, 2008; Kaufmann *et al.*, 2011; Apte *et al.*, 2017)

We derive a dissipative response time *τ* that measures the duration of viscoelastic transition towards equilibrium mechanics (Figs 3c-iii, 4c-iii, 5d-iii). Tethering of the nucleus to the surrounding cytoskeleton was shown to slow down strain relaxation and rate of mechanical energy release owing to cytoskeletal viscosity. (Wang *et al.*, 2018) The response time that we measure in live nematodes (minutes) is significantly longer than the equivalent time scales measured for isolated nuclei (< 1 second)(Wang *et al.*, 2018) and nuclei surrounded by cytoskeleton within intact cells (seconds).(Swift *et al.*, 2013b) Rapid release of mechanical energy increases the risk of structural damage to the nucleus and the encapsulated genetic material. We find here that the actual energy release rate within a live organism is even slower than *in vitro* estimates. Hence, the physiological environment of cells and tissues provide a dissipative mechanism for protecting the nucleus and chromatin from applied forces.

## Methods

### Non-aging strains and culture

We used the following *C. elegans* strains in this study: PD4810 (*lmn-1*:GFP, *ccIs4810* [pJKL380.4 P*lmn-1*::*lmn-1*::gfp::*lmn-1* 3’UTR + pMH86 *dpy20*(+)]), YG751 (*lmn-1*-GFP:*unc-84* null [pJKL380.4 Plmn-1::lmn-1::gfp::lmn-1 3’UTR + pMH86 dpy20(+);unc-84(e1174)]). *C. elegans* strains were maintained and manipulated under standard conditions as previously described.(Brenner, 1974) Strains were out-crossed at least 3 times to ensure a clean background.

### Nematode culture and stretching

*C. elegans* strains were bleached in 0.06 M NaOH, 0.5% sodium hypochlorite solution. L1 synchronization was performed by incubating the embryos on empty NGM plates for 18 hours at 20°C. L1 nematodes were washed in M9 buffer (3g KH2PO4, 6g Na2HPO4, 5g NaCl, 1ml 1M MgSO4, H2O to 1 liter sterilize by autoclaving) and reseeded in plates containing a thin lawn of OP50/RNAi bacteria. Late L4 synchronized nematodes were collected into Eppendorf tubes and washed 3 times. To support adhesion, 0.2% Poly-L-Lysine (Sigma P2636) was added at equal volume and supplemented with 10 μL anti-fade solution (9 parts glycerol, 1 part 10X PBS, 0.1 part 20%(w/v) n-propyl gallate (Sigma P3130) in dimethyl sulfoxide). Nematodes were allowed to settle down for a few minute.

Nematodes were transferred to the center of a 4″ × 4″ square silicon sheet (cut from 12″ × 12″, 0.005″ NRV G/G 40D silicon membrane, SMI) in twenty separate droplets. The nematodes were then covered by a second silicone sheet and trapped air bubbles were removed. The “sandwiched” nematodes were placed on a stretching device(Zuela *et al.*, 2016) and mounted on top of an inverted microscope (Nikon Eclipse Ti-U inverted microscope, equipped with a Nikon S Plan fluor ELWS 40X/0.60 for PD4810 experiments and a Plan Apo VC 20X/0.75 for YG751 experiments). Equiaxial stretching was performed within the conjugated plane of the microscope and time-lapse recording (Andor Zyla 4.2 sCMOS camera controlled with NIS Elements 4.5.00 software) was performed both in bright field phase contrast and fluorescence channels (Nikon Intensilight C-HGFI light source and Chroma ET - EGFP (FITC/Cy2) filter set).

### Image processing, mechanical and statistical analyses

Whole nematodes (bright field channel) and nuclei (EGFP fluorescence channel) were segmented using ImageJ. As described in the main text, projected area, contour length and distances were calculated using ImageJ tools. Viscoelastic analyses and curve fitting were performed as described in detail in the main text using Matlab-2018a (Mathworks).

### Cytoskeletal and nuclear drug perturbations

Trichostatin-A (TSA, Sigma T8852) was diluted in M9 medium at 2.5 mM. Five droplets (150 μl in total) were added onto the OP50 bacteria lawn of NGM plates to reach a complete coverage and left to dry. Similarly, Latrunculin-A (Lat-A, Sigma L5163) was freshly diluted in M9 medium at 5 mM. Five droplets (150 μl in total) were added onto the OP50 bacteria lawn of NGM plates to reach full coverage and left to dry. Synchronized L1 nematodes were placed on top of the drug-immersed plates at 20°C for 24 hours until reaching the appropriate developmental stage (late L4).

### RNA interference experiments

RNA interference (RNAi) knockdown of target genes was performed via bacteria-feeding as previously described.(Timmons *et al.*, 2001) E-coli clones were obtained from the Vidal’s and Ahringer’s RNAi libraries.(Kamath and Ahringer, 2003; Kim *et al.*, 2005) L4440 clone was used as an empty vector control. *Lmn-1* and *unc-84* knockdown was performed using the DY3.2 and F54B11.3 vectors, respectively. Specifically, synchronization L1 nematodes were transferred to feeding plates at 20°C and experiments were performed at late L4 stage as described above.

### Aging stretching experiments

Synchronized L1 PD4810 were placed on NGM plates at 20°C for 24 hours, until reaching late L4 stage. Late L4 nematodes (D0) were extracted by picking for subsequent experiments. The remaining nematodes were further incubated at 20°C for additional 12 hours until reaching adult stage. Adult nematodes were transferred by picking into new NGM plates and incubated at 20°C. Two-day old adult nematodes (D2) were extracted by picking 36 hours after transfer for subsequent experiments. The remaining nematodes were transferred by picking to new NGM plates and incubated for an additional 48 hours at 20°C. Four-day old adult nematodes (D4) were then extracted by picking for subsequent experiments. Extracted nematodes were transferred into M9 droplets and placed onto an unseeded NGM plate for 10 minutes to remove residual OP50 bacteria. The nematodes were then transferred into a single 1 μL droplet containing 3:1 M9:anti-fade mixture and placed at the center of a 4″ × 4″ square silicon sheet. Nematode stretching was performed by placing a second membrane on top and mounted onto the stretching apparatus as described above. Each age group consisted of 20 nematodes.

### Western Blotting

To eliminate the contribution of embryonic Ce-lamin to the calibration of adult Ce-lamin levels across age groups, we used temperature-sensitive sterile CF512 nematode strain [(b26) II; fem-1(hc17) IV]. Bleached CF512 embryos were synchronized as described above. L1 nematodes were transferred to OP50 containing NGM plates and incubated at 25°C for 24 hours. 150 late L4 nematodes (D0) were collected for Western blotting and the remaining nematodes were transferred to new NGM plates and incubated at 25°C (24 hours) followed by 20°C (24 hours). 150 two-days old (D2) nematodes were collected for Western blotting. The remaining nematodes were transferred to new NGM plates and incubated at 20°C (48 hours). 150 four-days old (D4) nematodes were collected for Western blotting.

Western blotting was performed as previously described.(Towbin *et al.*, 1979; Bank *et al.*, 2011) In short, nematode samples were washed three times in M9 buffer and immersed in 100 μL M9 buffer. Nematodes were lyzsed by adding 900 μL mixture of RIPA lysis Buffer (10mM Tris-Cl pH 8.0, 1mM EDTA, 0.5mM EGTA, 1% Triton X-100, 0.1% Sodium Deoxycholate, 0.1% SDS, 140mM NaCl), X25 protease inhibitor (cOmplete protease inhibitor cocktail by Sigma Aldrich) and 0.1M PMSF (Sigma P7626) at 95:4:1 volume ratios. Nematode lysate solutions were centrifuged at 2,000 rpm for two minutes. Supernatant was removed and pellet was dissolved in 200 μL RIPA-PI-PMSF lysis buffer. The samples underwent four freezing-thawing cycles in liquid nitrogen and sonication (Sonics Vibra Cell sonicator with a model CV33 microtip at 34% amplification 4×10 sec sonication periods). Samples underwent two freeze-thaw cycles and centrifuged 7,500 rpm for 5 minutes at 4°C. Supernatant was collected and transferred to a new Eppendorf tube for use.

Total protein levels were calibrated using Bicinchoninic Acid (BCA) assay (ThermoFisher Pierce BCA Protein Assay kit, OD measured with BioTek synergy 2 multi-mode multiplate reader). Samples were ran on a 9% SDS polyacrylamide gel, transferred to a nitrocellulose membrane and blotted using anti Ce-lamin antibody (KLH-conjugated C-terminal peptide of Ce-lamin; VEFSESSDPSDPADRC; serum 3932, bleed 6, dilution of 1:1000). Chemiluminescence images were acquired (Vilber Fusion FX). Ce-lamin levels were quantified by measuring total band intensities normalized to total protein levels.

## Supplementary material

**Figure-S1.**
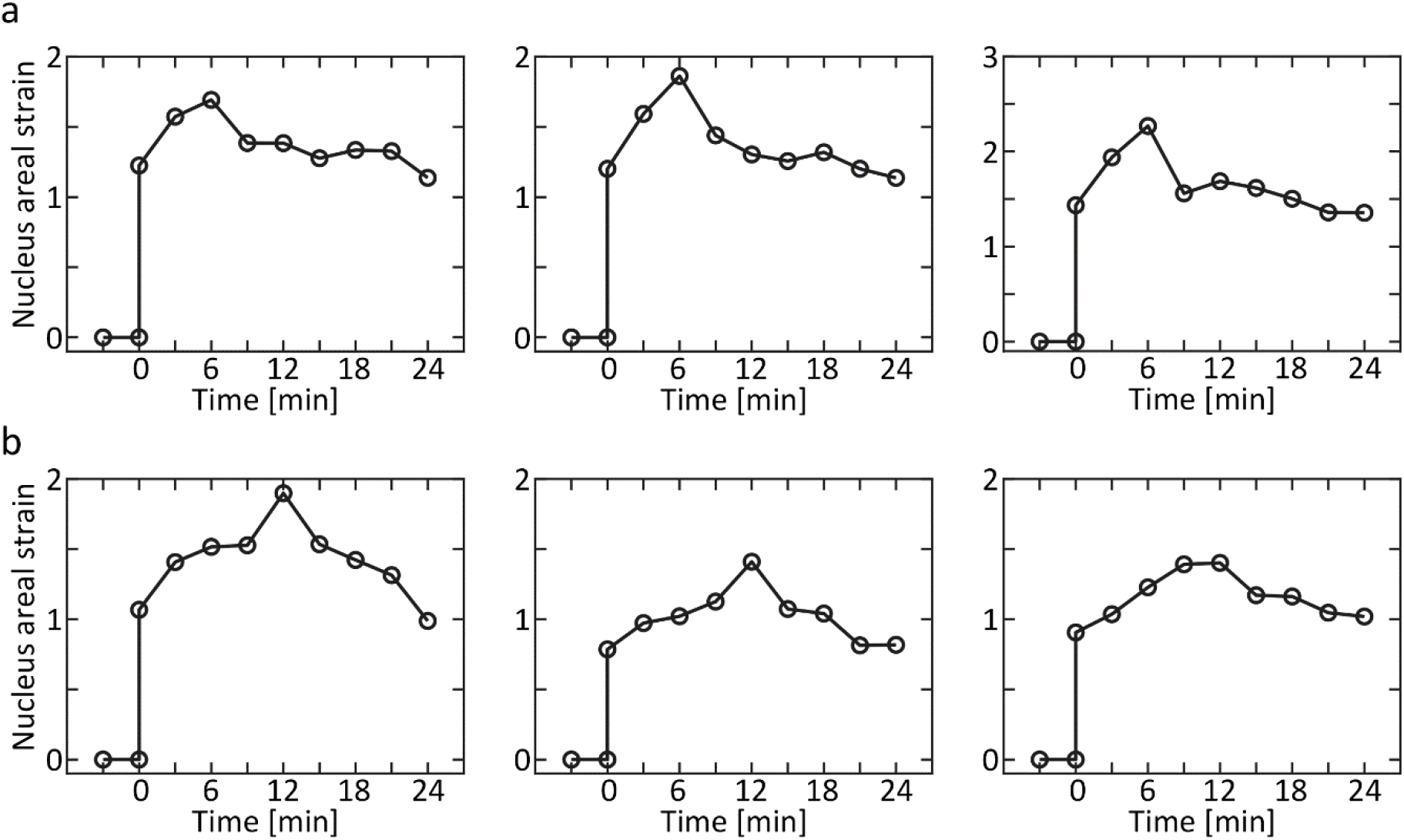
Individual nuclei respond non-monotonically to constant load. Dynamic areal strain profiles of three representative nuclei within (a) muscle and (b) hypodermis tissues are depicted. The non-monotonic mechanical response shared by all nuclei in both tissues is characterized by an instantaneous elastic deformation, creep and deformation recovery.

