## Supplementary figure S1 for "Measuring nucleus mechanics within a living multicellular organism: Physical decoupling and attenuated recovery rate are physiological protective mechanisms of the cell nucleus under high mechanical load"

### Supplementary material 1

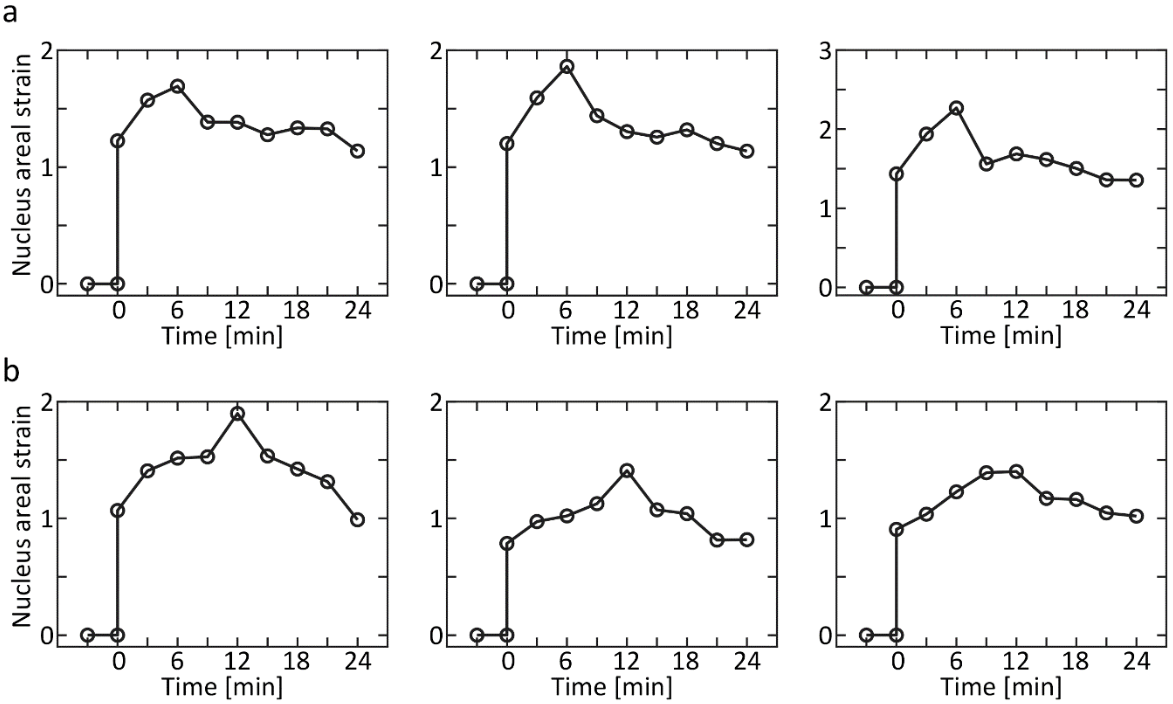


**Figure-S1 | Individual nuclei respond non-monotonically to constant load.** Dynamic areal strain profiles of three representative nuclei within (a) muscle and (b) hypodermis tissues are depicted. The non-monotonic mechanical response shared by all nuclei in both tissues is characterized by an instantaneous elastic deformation, creep and deformation recovery.

2
